# Quantitative comparison of a mobile, tablet-based and two stationary, video-based eye-trackers

**DOI:** 10.1101/2023.06.12.544554

**Authors:** Aylin König, Frank Bremmer, Stefan Dowiasch

**Author notes:** shared senior-authorship (Frank Bremmer, Stefan Dowiasch). Corresponding author: Dr. Stefan Dowiasch, Philipps-Universität Marburg Fachbereich Physik, AG Neurophysik, Karl-von-Frisch-Straße 8a 35043 Marburg / Lahnberge.

## Abstract

The analysis of eye-movements is a non-invasive, reliable and fast method to detect and quantify brain (dys-)functions. Here, we investigated the performance of two novel eye-trackers: the Thomas-Oculus-Motus-research mobile (TOM-rm) and the TOM-research stationary (TOM-rs) and compared it with the performance of a well-established video-based eye-tracker, i.e., the EyeLink 1000 (EL). The TOM-rm is a fully integrated, tablet-based mobile device that presents visual stimuli and records head-unrestrained eye-movements at 30Hz without additional infrared (IR) illumination. The TOM-rs is a stationary, video-based eye-tracker that records eye-movements at either high spatial or high temporal resolution. We compared the performance of all three eye-trackers in two different behavioral tasks: pro– and anti-saccade and free viewing. We collected data from human subjects while running all three eye-tracking devices in parallel. Parameters requiring a high spatial or temporal resolution (e.g., saccade latency or gain), as derived from the data, differed significantly between the EL and the TOM-rm in the pro– and anti-saccade task. In the free viewing task, larger noise and the lower frame rate of the TOM-rm caused deviations of the results with respect to the EL. Differences between results derived from the TOM-rs and the EL were most likely due to experimental conditions, which could not be optimized for both systems simultaneously. We conclude that the TOM-rm can be used for measuring eye-movements reliably at comparably low spatial and temporal resolution. The TOM-rs, on the other hand, can provide high-resolution oculomotor data at least on a par with an established reference system.

## Introduction

Eye-movements are a window to the mind and brain (van Gompel et al., 2007). They offer a reliable and non-invasive way to detect and quantify brain functions and their neural correlates (Leigh & Kennard, 2004; Coe & Munoz, 2017). As documented already more than a century ago, disorders of eye-movements are often related to neurological and psychiatric diseases, even already at an early stage of the disease (Diefendorf & Dodge, 1908). There is a variety of eye-tracking methods, ranging from electro-oculogram (EOG) via the search-coil-method to video-oculography. Even for video-based eye-trackers there are huge differences in application areas and performance (and costs). Accordingly, it is essential that a potential user is provided with all relevant information about the performance and usability of the eye-tracker (Hutton, 2019). In this study, we investigated the performance and usability of two new eye-trackers, which were developed with the long-term goal of serving as a support tool in diagnostics for neurological and psychiatric diseases. The *Thomas-Oculus-Motus – research mobile (TOM-rm)* is a tablet-based device, which can be used in an everyday setting (e.g., at home) without additional infrared illumination. The *TOM – research stationary (TOM-rs)* is a video-based eye-tracker meant for high-resolution recordings in a lab environment. To investigate the performance and usability of both TOM eye-trackers, we measured a set of eye-movements in healthy participants concurrently with a third eye-tracker, i.e., the EyeLink 1000 (EL, SR Research), a video-based eye-tracker, which is well established in oculomotor research. We focused our study on eye-movements and related functions that have been shown to be compromised in neurological and psychiatric diseases, i.e., (pro– and anti-)saccades (Antoniades et al., 2015; Coe & Munoz, 2017; Leigh & Zee, 2015), free viewing (Matsumoto et al., 2011), and dynamics of pupil responses (Wang et al., 2016).

## Methods

### Participants

A total of 30 subjects participated in the pro– and anti-saccade task (i.e., a saccade in the opposite direction with respect to the target), 16 male and 14 females with a mean age of 24.86 ± 3.50 years. 29 of the 30 subjects who participated in the pro– and anti-saccades task, participated also in the free viewing task, 15 male and 14 females with a mean age of 24.90 ± 4.07 years. Inclusion criteria were: (i) no glasses, (ii) no color weakness, and (iii) no history of neuropsychiatric impairments. In oculomotor studies, typically the performance of the participants is monitored online and sometimes trials have to be repeated because a participant’s gaze left an invisible control window. Yet, the TOM-rm did not allow for such an online control during data recording, since it first records videos of the participants’ eyes followed by an automatic extraction of eye traces in an offline analysis. Importantly, we measured subjects’ oculomotor performance with all three systems at the same time. This approach has the advantage that comparisons between systems can be made on a trial-by-trial basis. The downside of this approach, however, is that the experimental conditions could not be optimized for all three eye-tracking systems simultaneously. This concerns especially the infrared illumination for the two high-speed video-based eye trackers, but also missed trials due to the non-existent online gaze control of the TOM-rm. Accordingly, offline analysis revealed that not all data from all participants could be considered in our population analysis. Consequently, data from nine subjects involved in the pro– and anti-saccade task and data from eight other subjects participating in the free viewing task had to be excluded from further analysis since we couldn’t track their pupils (and hence eye position) reliably throughout the recording with all three eye-trackers simultaneously.

Subjects were remunerated with 8€/hour. All procedures used in this study were in accordance with the Declaration of Helsinki and were approved by the local ethics committee (AZ-2012-23K).

### Eye-tracker and laboratory setup

The laboratory setup for concurrent data collection with all three eye-trackers is shown in Figure 1.

**Figure 1:**
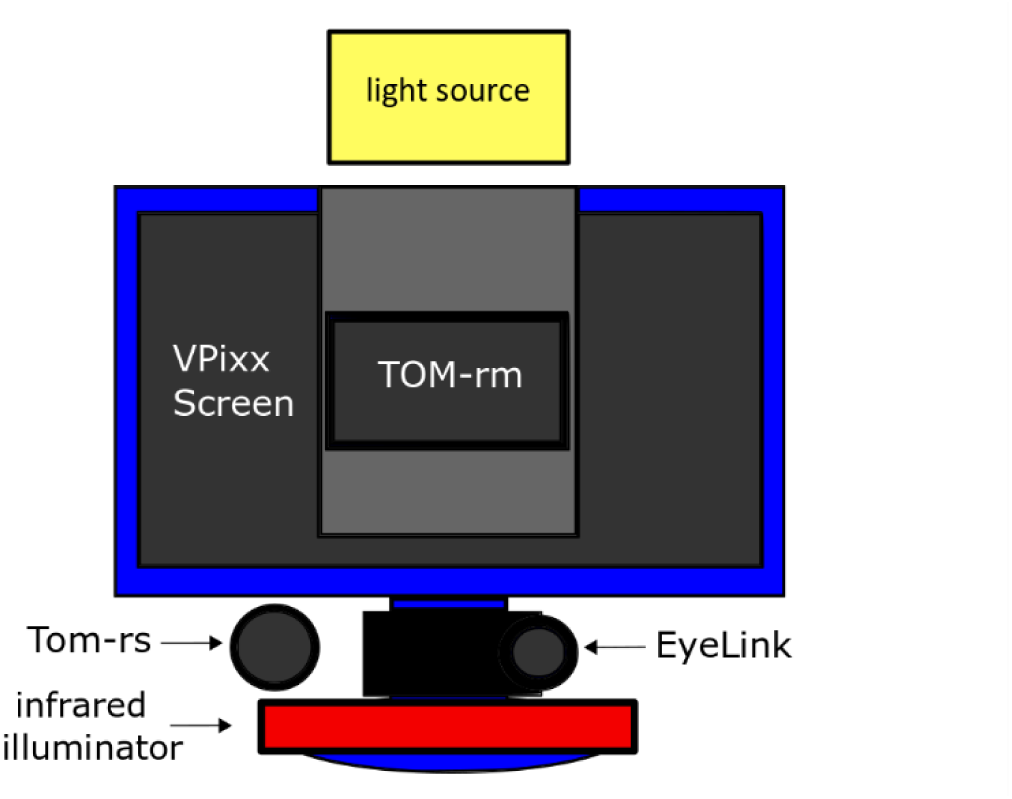
Setup to measure with all three eye-trackers in parallel. The VPixx monitor was placed centrally in front of the subject. The TOM-rm was positioned in its customized holder exactly in the center on the VPixx monitor. The TOM-rs was placed to the left and below the screen. The EyeLink 1000 was positioned directly below the screen.

The experiment took place in a soundproof room with indirect lighting, which was necessary because the TOM-rm only detects the gaze position with visible light. Subjects sat in front of a screen at a viewing distance of 45cm with their head stabilized by a chin-rest and a forehead support. Visual stimuli were presented binocularly on the display of the TOM-rm (Huawei MediaPad M5, height 11.4cm, width 18cm) at a resolution of 2560 x 1600px and a framerate of 30Hz. Simultaneously, a video of the participant’s head was recorded with the tablet’s front camera (8MPx) at a framerate of 30Hz. Under optimal experimental conditions, the TOM-rs records eye-movements either with high spatial (e.g., Full-HD @ 150Hz) or high temporal (e.g., 640 x 480px @ 2000Hz) resolution. Here, we measured with a frequency of 500Hz and a resolution of 640 x 480px. This relatively low frame rate was necessary to compensate for the lower IR light sensitivity required by the EL. The TOM-rs camera was placed to the left and below the participant’s head next to the centrally positioned EL. Both, the TOM-rs and the EL, need an individually adjusted IR illumination. The EL requires a less intense IR light-source, because it uses a lens with a fixed focal length but high light intensity. The TOM-rs on the other hand uses a zoom lens with a variable focal length between 16mm-300mm, and hence requires IR-illumination at a higher intensity. To be able to measure with all eye-trackers simultaneously, we had to adjust both IR illuminators (EL and TOM-rs) such that both eye-tracking systems could reliably detect the participants’ pupil. The EL-system was set to an intensity of 75% of its infrared illumination to avoid over-illumination by the additional IR light source of the TOM-rs. The infrared intensity of the TOM-rs illuminator was adjusted by modifying the angle of the irradiated infrared light in a subject-specific manner. This necessary tradeoff, however, has the potential to compromise overall data-quality due to non-optimal lighting conditions (see also Discussion).

### Procedure of the experiment

First, we calibrated the EL with a nine-point calibration task presented on the standard lab monitor (VPixx / 3D Lite, 1920 x 1080 pixels @ 120 Hz). Next, we calibrated the TOM-eye-trackers, with a nine-point calibration stimulus displayed on the TOM-rm screen. The fixation points appeared in a pseudo randomized order for two seconds each, with an amplitude of ±10.8° in the horizontal direction and ±6.7° in the vertical direction with respect to straight-ahead. In the following, all experimental stimuli (pro– and anti-saccade; free-viewing) were presented on the screen of the TOM-rm.

### Pro– and anti-saccades task

In this task, 40 pro– and 40 anti-saccade trials were presented in pseudo-randomized order (Figure 2a). At the beginning of each trial, a blue or red fixation point (FP) with a diameter of 1° visual angle was displayed on a grey background in the center of the screen. The participants were asked to fixate this target. The color of the fixation target (blue or red) indicated whether a pro– or anti-saccade should be performed during the further course of the trial (blue: pro-; red: anti-saccade). After 1s the FP was switched off. 200ms later, a white target point (TP) with a diameter of 1° visual angle appeared for one second in pseudo-randomized order 10.8° to the right or left of the FP on the horizontal meridian (HM). The subjects were instructed to perform the pro– or anti-saccade as quickly as possible. Between trials, a grey screen was presented for 1 second. In total, this paradigm took about five minutes.

**Figure 2:**
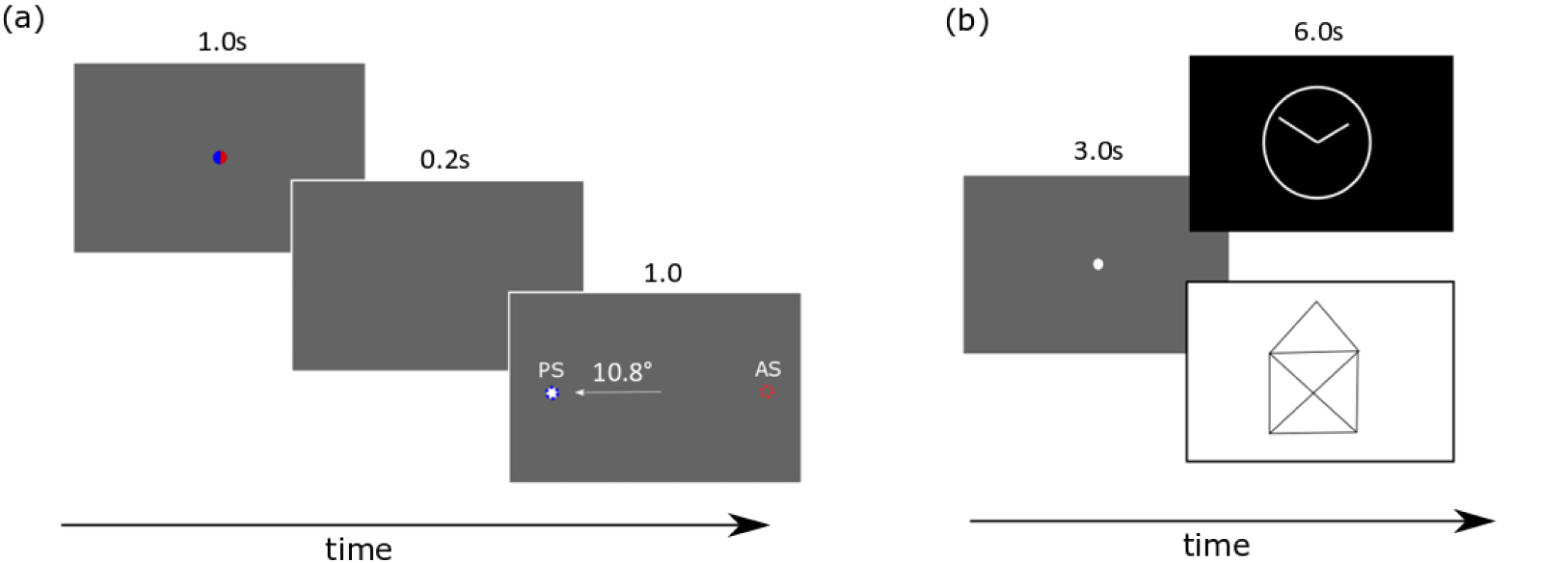
(a) Schematic illustration of the pro– and anti-saccades task. A blue or a red fixation point (FP) appeared for 1s in the center of the screen, signaling the subject to perform a pro-saccade (blue FP) or anti-saccade (red FP). After a gap of 0.2s, a white target point with a diameter of 1° visual angle appeared for one second in pseudo-randomized order 10.8° to the right or left of the FP. (b) Schematic illustration of the free viewing task. After a three second fixation period a light (as shown on the bottom right) or a dark (as shown on the top right) geometrical image appeared for six seconds. The subjects were instructed to look closely at the pictures. 30 images (15 bright and 15 dark) were shown in pseudo randomized order, while bright and dark pictures alternated.

### Free Viewing Task

In the second task, static images were displayed on the screen for a duration of 6s (Figure 2b).

We chose simple line drawings like a house for this task, because it has already been shown that the visual exploration of images like these is impaired in neuropsychiatric diseases like Parkinson’s disease (Matsumoto et al., 2011; Archibald et al., 2013). Before each image was presented, a central white fixation point with a diameter of 1° visual angle was displayed on a grey background for 3 seconds. This period served as an individual reference for the analysis of the change in pupil diameter after the much brighter or darker images were presented. Participants were instructed to fixate the initial fixation target and then were allowed to freely move their eyes and inspect the images for six seconds. A total of 30 images (15 bright images, i.e., black lines on a white background and 15 dark images, i.e., white lines on a black background) were shown, resulting in a total duration of this measurement of approximately 5 minutes.

## Data analyses

Data analysis was performed using MATLAB R2018a (The MathWorks Inc.). Typically, different eye-trackers come with different software packages for analyzing eye-movements, which could have an influence on some eye-movement parameters such as saccade mean velocity (Dowiasch, Wolf, & Bremmer, 2020). Accordingly, as a first step, we analyzed our data with the software provided with the eye-tracker (in case of the EL) or which we developed specifically for the eye-tracker (TOM-rs and –rm). In the following we call this approach *individual evaluation*. In the case of the EL the detection of saccades and fixations was provided by the EL data analysis package. In case of the TOM-eye–trackers, we used a custom build software for the detection of saccades and the EyeMMV toolbox for detecting fixations (Eye Movements Metrics & Visualizations; National Technical University of Athens) (Krassanakis, Filippakopoulou, & Nakos, 2014). In a second step, we used one common toolbox (developed for the TOM-rm) to analyze the raw data of all three eye-tracking systems to allow for a quantitative comparison of their performance. We refer to this evaluation method as *same evaluation*. To this end, data from the TOM-rs and the EL were down-sampled to 30 Hz, i.e., the genuine sample rate of the TOM-rm, by linear interpolation. Due to the different internal clocks of the three eye tracking systems, we had to synchronize their times series data. These eye-tracking data were synchronized in time by cross-correlation. To this end, we correlated the horizontal eye position data of the EL and the TOM-rs with those of the TOM-rm. This procedure allowed us to shift eye position traces in time such that the EL and TOM-rs data were matched in time to the TOM-rm data.

### Saccades

In the *individual evaluation*, saccades from the EL data were determined using the saccade detector provided by the manufacturer, which is based on a combined saccade-related position-, velocity-, and acceleration-threshold (0.1° to 0.2°, 30°/s, 8000°/s^2^). Furthermore, we developed two different, but similar saccade detectors for the two TOM– eye-trackers. The saccade detector of the TOM-rm aims for data with a low sample-rate and employs a pure velocity criterion of 20°/s. Furthermore, the duration of a saccade must not exceed 200ms. Before determining the velocity of the eyes, the eye position data were filtered using a median filter over 150ms in order to minimize noise and at the same time not reduce the saccade velocity (Juhola, 1991). For the saccade detector of the TOM-rs we filtered the velocity with a second-order Savitzky-Golay filter over 22ms (Savitzky & Golay, 1964). For detection of saccade on– and offsets we used a velocity criterion of 50°/s. Also, for the saccade detection with the TOM-rs, the duration of a saccade must not exceed 200ms. For the pro– and anti-saccade tasks, in order to focus analysis on the task specific saccades and exclude small corrective saccades, only saccades larger than 3° and smaller than 20° were considered. For determining the error rate of pro– and anti-saccades only those trials were considered for which the saccade start did not deviate from the central fixation spot by more than ±2.5 degrees in the horizontal direction. All trials with a saccade latency between 100ms and 450ms were considered as valid (Wang et al., 2016; Waldthaler et al., 2021). Only those trials were evaluated in the pro– and anti-saccades task which fulfilled the previously mentioned criteria by all three eye-trackers. This rather harsh but necessary criterion resulted in a large number of dropouts. Furthermore, it could happen that different trials were approved as valid for the *individual* and *same evaluation*. For example, in a certain trial, no saccade was detected in one eye-tracker during the *same evaluation*. In the *individual evaluation*, however, a saccade was detected in exactly this trial in all three eye-trackers.

Accordingly, the approach led to an unequal number of trials in the two data analysis procedures. In total participants performed 1646 trials, where one subject only completed 46 trials instead of 80 trials. Under the previously mentioned criteria, 1272 trials in the *individual evaluation* and 1233 trials in the *same evaluation* were evaluated as valid by the EL, 1185 trials in the *individual evaluation* and 1189 trials in the *same evaluation* by the TOM-rs, and 1100 trials in the *individual evaluation* and in the *same evaluation* by the TOM-rm (Table 1).

**Table 1:**
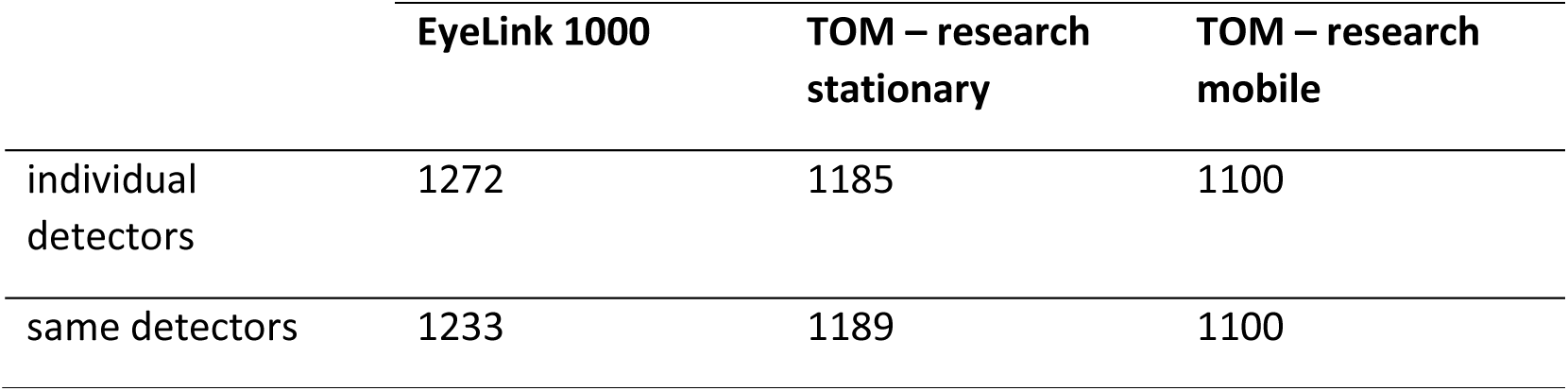
Number of valid trials for the EyeLink 1000, the TOM – research stationary and the TOM – research mobile. Evaluation with the *individual* and *same evaluation* analysis algorithms.

|  | EyeLink 1000 | TOM – research stationary | TOM – research mobile |
| --- | --- | --- | --- |
| individual detectors | 1272 | 1185 | 1100 |
| same detectors | 1233 | 1189 | 1100 |

In total 1024 trials were detected with all three eye-trackers in the *individual evaluation* and 999 trials in the *same evaluation* and were included in the further analysis. Another parameter we determined was the yield, which we defined as the ratio of the number of detected saccades and twice the number of valid trials.

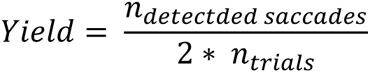

In the ideal case, the yield should be 1.0, since each trial should trigger two saccades, i.e., the saccade towards the saccade goal and a saccade back to the central fixation. Yet, since there were trials in which correction saccades occurred, or in which subjects did not return their gaze back to the central fixation with a single saccade, or in which noise was detected as a saccade, often more than two saccades were detected per trial, possibly resulting in a yield > 1.

In the free viewing task only saccades larger than 1° amplitude were included for all three eye-tracker, because saccades smaller than 1° could not always be reliable detected with the TOM-rm.

### Fixations

For the *individual evaluation* of the data, the detection of the fixations for the EL was performed with the built-in detector from the manufacturer. Here, all eye traces that do not qualify as a saccade or smooth pursuit are considered a fixation. The detection of fixations for the TOM-eye-trackers was performed using the EyeMMV toolbox. The lower limit of the fixation time for both TOM-eye-trackers was set to 150ms. In addition, in this toolbox, two spatial thresholds t1 and t2 have to be defined, which determine the maximum eye position jitter allowed to consider an eye trace as fixation (Krassanakis et al., 2014). We used values of t1 = 1° and t2 = 0.7° for evaluation with the TOM-rm data and t1 = 0.5° and t2 = 0.3° for evaluation of the TOM-rs data. For the *same evaluation* applied to data from all three eye-trackers, we used the fixation detector of the EyeMMV toolbox with the thresholds t1 and t2 of the TOM-rm. Before, the eye-position data of the pro– and anti-saccade task of the TOM-rm were filtered using a median filter over 0.3s, since they are subject to high noise. The TOM-rs eye-position data were also filtered over 0.2s due to the noise resulting from the suboptimal illumination condition. We also filtered the eye-position of the TOM-eye-tracker of the free viewing task with a median filter (TOM-rm & TOM-rs *same evaluation*: 0.2s; TOM-rs *individual evaluation*: 0.05s).

### Pupillary response

The pupillary response was only recorded with the EL and the TOM-rs systems. The light and dark reflex of the pupil were determined in the free viewing task during the observation of the light and dark images. The light reflex is typically considered the beginning of the constriction of the pupil during viewing bright images. In contrast, the dark reflex is considered the beginning of the pupil dilation while viewing dark images. In line with published work (Bergamin & Kardon, 2003) we defined the onset of the light reflex as the point in time when the acceleration of the value of the pupil area was maximally negative. Likewise, the onset of the pupil dilation was defined as point in time when this acceleration was maximally positive. For the determination of the change in pupil size we determined the z-score of the given pupil area, and we set the change in pupil area to 0 arbitrary units (a.u.) at time t = 0s. Further important parameters for the determination of the pupil dynamics are the minimum area during maximum constriction and the time this minimum area is achieved during the observation of the light images (see Figure 3).

**Figure 3:**
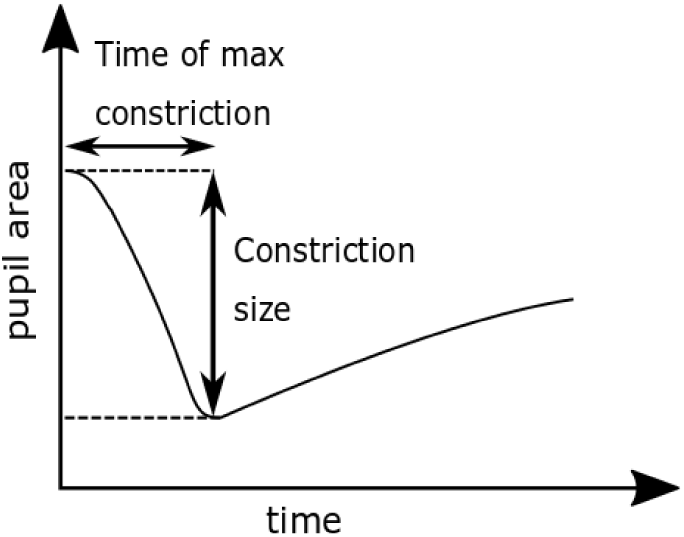
Schematic representation of the determination of the two parameters constriction size and time of maximum constriction.

We did not calculate these two parameters for the dilation, because due to the slow dark adaptation the average pupil area was about to reach a maximum value only at the end of a given trial (Wang & Munoz, 2014). We also aimed to quantitatively compare the dynamics of the pupil area as measured from the stationary eye-trackers. Therefore, we averaged the size of the pupil area over all subjects and determined the 95% confidence interval, separately for bright and dark images.

### Blink detection

Blink’s artifacts and all data points ±60ms around a blink were excluded from further analysis. For the *individual evaluation* of the EL data we employed the built-in eyelid blink detector, where blinks are defined as periods of data for which the pupil cannot be detected. For the TOM-rs we first applied a median filter (width: 115samples) on the pupil area data. All samples of this new time series data that deviated by more than a specific value, which we determined for every participant’s dataset individually, were considered blinks. In the same way, we detected the blinks of the EL for the *same evaluation*. Due to the fact that the TOM-rm does not make use of IR-illumination, no pupillometry data were available for the TOM-rm, but the Eye-Aspect-Ratio (EAR) (Cech & Soukupova, 2016). With the EAR, blinks were detected in the same way as with the TOM-rs, but with a larger width of the samples of the moving average (width: 35 samples, corresponding to > 1s). This long interval was necessary because of the relatively high noise in the EAR data. Accordingly, only a rough measure of the EAR was created. Both the blinks in the pupil area data and in the EAR were expressed by short, extreme peaks in the time series data.

Finally, saccades were only included in our data analysis if no blink was detected from 200ms before until 200ms after the saccade.

### Statistics

For the statistical analysis, we performed a paired-sample t-test to probe for differences in the quantitative eye movement parameters derived from data recorded with the three eye-trackers. Differences were considered significant if the p-value was smaller than α = 0.05. We also performed a Bonferroni correction, because for a given dataset, we performed more than one statistical test. This is why p-values had to be corrected for multiple comparisons by multiplying the uncorrected values with the number of hypotheses (pro-/anti-saccade task: 6 hypotheses, free viewing task: 3 hypotheses), resulting in new p-values. Consequently, this approach allows us to keep the typical alpha level, indicating significant differences, at 0.05.

In this study we used a rather unconventional method to determine the standard deviation (SD) in order to probe for differences in eye position data between the eye-trackers. We used this method detailed below because the *inter-individual* variance, i.e., variance of data across subjects, could potentially mask the differences in the data caused by the eye-trackers. We determined the SD as follows: First, for each subject, the mean value across all three eye-trackers of a given parameter 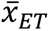 was determined. I.e., if the saccade latency for one subject is *x*_*EL*_ for the EL, 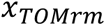 for the TOM-rm and 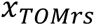 for the TOM-rs, then we determined the mean value 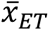 as follows:

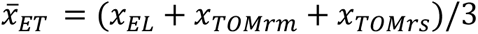

The mean value per subject per eye-tracker *x*_*EL*_, 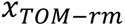 and 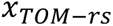 was subtracted from the mean value over all eye-trackers 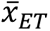

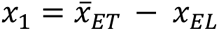

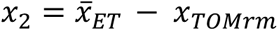

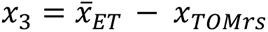

Then the SD was calculated over these values. In the following we called this the *inter-device SD*.

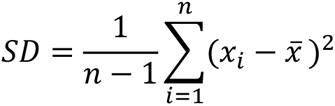

*x_i_* is the i-th value in the data set, 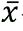 is the mean value of the data set and n is the sample size.

## Results

Here, we compared the performance of two novel eye-trackers, the TOM-rm and the TOM-rs, with a well-established eye-tracker in the field, the EyeLink 1000 (EL). Typical eye movement parameters were analyzed in two different behavioral tasks (pro– and anti-saccade and free viewing). In a first step, we analyzed data from each eye-tracker individually, i.e., with software either provided by the manufacturer (EL) or customized to the specifics of the eye-tracker (TOM-rm and TOM-rs), called *individual evaluation* in the following. In a second step, we analyzed data from all three eye-trackers with the same data analysis programs, i.e., those developed for the TOM-rm. We refer to this approach as *same evaluation* in the following. All mean values, inter-device SD and inter-individual SD of the pro– and anti-saccade task can be obtained from the supplements in Table S1. The respective results of the free viewing task are provided in Table S2.

### Pro– and anti-saccade task

The error rate of pro-saccades (t_rm_(20) = –1.40, t_rs_(20) = –1.40, p_rm_ = 1.00, p_rs_ = 1.00) and anti-saccades (t_rm_(20) = –1.38, t_rs_(20) = –1.92, p_rm_ = 1.00, p_rs_ = 0.42) as determined with the *individual evaluation* were not significantly different between the data obtained with the EL and the TOM-rm (as indicated by p_rm_ values) or the EL and the TOM-rs (as indicated by the p_rs_ values) (Figures 4a,b). Only correct pro– and anti-saccades were used for computing the saccadic gain. We observed a significant difference between data from the EL and TOM-rm concerning the gain in the pro– and anti-saccade task (pro: t_rm_(20) = 6.20, p_rm_ = 2.79 x 10^-5^, anti: t_rm_(20) = 7.40, p_rm_ = 2.30 x 10^-6^). On average, the gain as determined from the TOM-rm data was 7% smaller as compared to the gain derived from the EL-dataset. There were no such differences between data obtained with the EL and the TOM-rs (pro: t_rs_(20) = 1.30, p_rs_ = 1.00, anti: t_rs_(20) = 1.56, p_rs_ = 0.81. Figure 4c). For the pro-saccade latency, we found a significant difference between the EL and the TOM-rm datasets, but not between the EL and the TOM-rs data-sets (t_rm_(20) = 13.20, t_rs_(20) = –1.43, p_rm_ = 1.49 x 10^-10^, p_rs_ = 1.00). This was also the case for the anti-saccade latency (t_rm_(20) = 10.75, t_rs_(20) = –2.59, p_rm_ = 5.53 x 10^-9^, p_rs_ = 0.10. Figure 4d). In both cases, pro– and anti-saccades, saccadic latencies derived from the TOM-rm dataset were about 18ms shorter than those values determined from the EL-dataset. Figure 4e shows the average number of saccades. We found no significant differences between results from the eye-trackers concerning the number of saccades (t_rm_(20) = 2.91, t_rs_(20) = 0.78, p_rm_ = 0.05, p_rs_ = 1.00). In the ideal case, participants would have performed 160 saccades each (80 trials, one saccade towards the peripheral stimulus and one back to the center of the screen). To have a reference for the number of saccades, we determined the yield, i.e., the ratio of the number of detected saccades and twice the number of valid trials. No significant difference between the eye-trackers could be determined concerning the yield. We found the following values: yield _EL_: 0.86 ± 0.07; yield _TOM-rm_: 0.97 ± 0.14; yield _TOM-rs_: 0.92 ± 0.11; t_rm_(20) = –2.67, t_rs_(20) = –2.18, p_rm_ = 0.08, p_rs_ = 0.25). Finally, we observed no significant difference between the data obtained with the EL and the TOM-eye-trackers concerning mean fixation durations for peripheral targets (t_rm_(20) = –0.21, t_rs_(20) = –0.12; p_rm_ = 1.00, p_rs_ = 1.00. Figure 4f).

**Figure 4:**
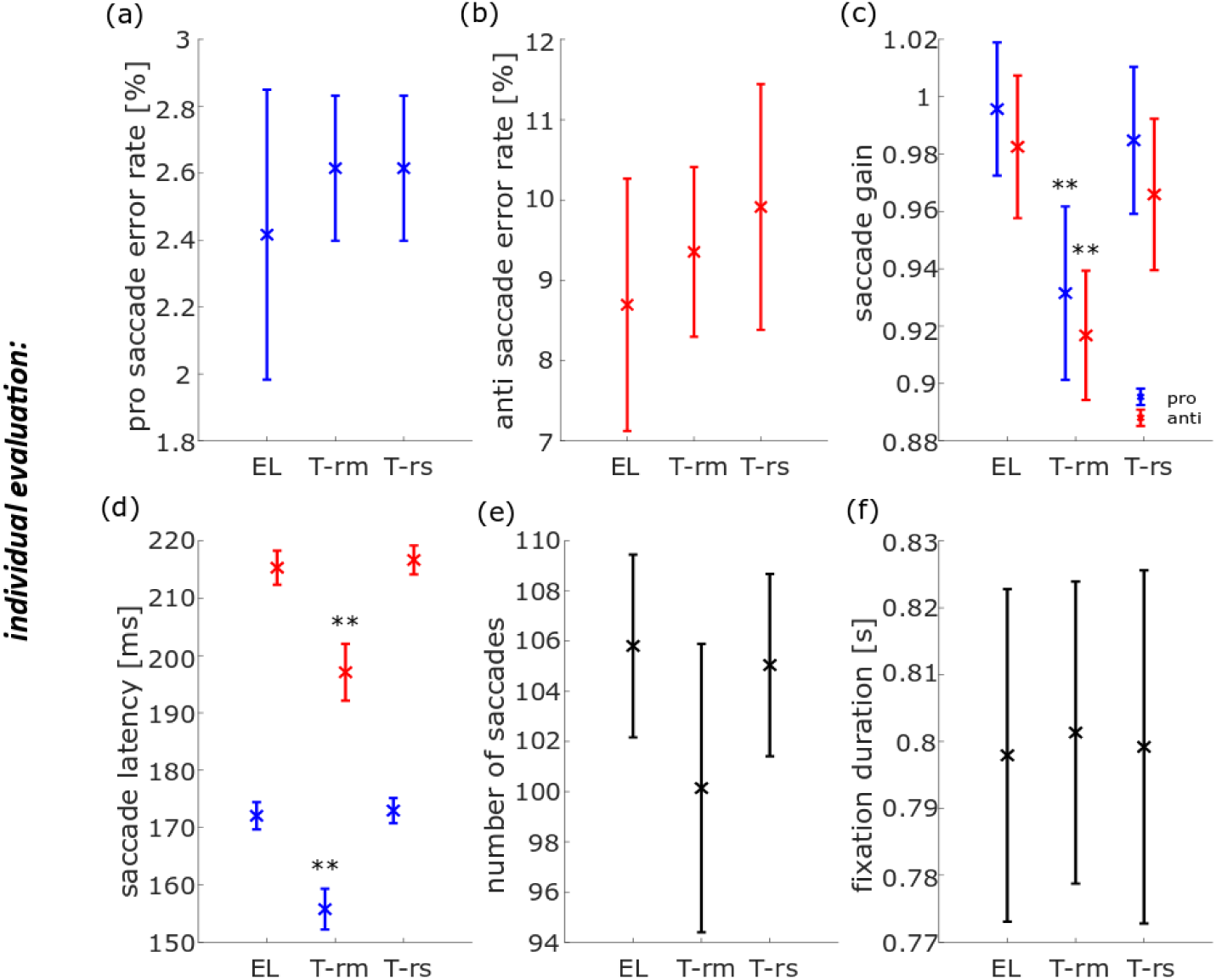
Population data for six eye movement parameters in the pro– and anti-saccade task, measured with the EyeLink 1000 (EL), TOM-research mobile (TOM-rm) and TOM-research stationary (TOM-rs) and analyzed with individual saccades and fixation detectors. The blue data points and their inter-device SD represent pro-saccade data, the red points and their inter-device SD the anti-saccade data and the black points and their inter-device SD pro– and anti-saccade data. (a) The error rate of pro-saccades, (b) error rate of anti-saccades, (c) mean saccade gain, (d) mean saccade latency, (e) mean number of saccades (f) and mean fixation duration. *p < 0.05, **p < 0.01.

In a second step, we compared the performance of the eye-trackers by analyzing the data with the *same evaluation* approach. For this purpose, we used the routines as developed for analyzing the TOM-rm data, since this eye-tracker provides data with the lowest spatial and temporal resolution. To make results comparable, we first down-sampled EL and TOM-rs data to 30Hz, i.e., the genuine sample rate of the TOM-rm (see Methods for details). No difference was found for the pro-saccade error rates neither between the EL and TOM-rm datasets nor between the EL and TOM-rs datasets (Figure 5a). Likewise, we did not find significant differences for the error rates in the anti-saccade task for the TOM-rm (t_rm_(20) = 1.27, p_rm_ = 1.00. Figure 5b). The anti-saccade error rate of the EL and the TOM-rs were the same. There was a significant difference of about 3%, though, between the EL and the TOM-eye-trackers in the pro– and anti-saccade gain (pro: t_rm_(20) = 9.29, t_rs_(20) = 3.97, p_rm_ = 6.43 x 10^-8^, p_rs_ = 4.55 x 10^-3^; anti: t_rm_(20) = 6.89, t_rs_(20) = 4.00, p_rm_ = 6.46 x 10^-6^, p_rs_ = 4.20 x 10^-3^; Figure 5c). Also here, the gain as determined from the TOM-rm data was about 7% smaller than its corresponding value determined from the EL-dataset. Unlike in the *individual evaluation*, a significant difference (< 3 ms) in pro– and anti-saccade latency between the EL and the TOM-rs datasets could be determined for the pro-saccade latency, but not for the anti-saccade latency (pro: t_rs_(20) = 4.17, p_rs_ = 2.84 x 10^-3^, anti: t_rs_(20) = 2.67, p_rs_ = 0.09). Confirming the results from the *individual evaluation*, also with the *same evaluation* we found a significant difference between the EL and the TOM-rm datasets, with latencies derived from the TOM-rm being on average about 13ms shorter than those derived from the EL-dataset (pro: t_rm_(20) = 8.00, p_rm_ = 7.05 x 10^-7^, anti: t_rm_(20) = 8.24, p_rm_ = 4.41 x 10^-7^; Figure 5d). The number of saccades as determined with the *same* saccade detector for the EL and the TOM-rm datasets were significantly different: roughly 109 as derived from the EL-dataset and 103 as derived from the TOM-rm dataset. There was no such difference between the dataset of the two stationary eye-trackers (t_rm_(20) = 3.91, t_rs_(20) = –0.88, p_rm_ = 5.15 x 10^-3^, p_rs_ = 1.00. Figure 5e). The yield of the number of saccades for the EL was 0.92 ± 0.08, for the TOM-rm 0.98 ± 0.11 and for the TOM-rs 0.97 ± 0.11. These values were not significantly different (t_rm_(20) = –1.98, t_rs_(20) = –1.49, p_rm_ = 0.37, p_rs_ = 0.91). Furthermore, we found a significant difference between the EL and the TOM-rm dataset concerning the mean fixation duration for the peripheral targets (0.93s for the EL-dataset and 0.80s for the TOM-rm dataset). However, we found no significant difference between the two stationary eye-tracker datasets (t_rm_(20) = 6.70, t_rs_(20) = 1.49, p_rm_ = 9.65 x 10^-6^, p_rs_ = 0.90. Figure 5f).

**Figure 5:**
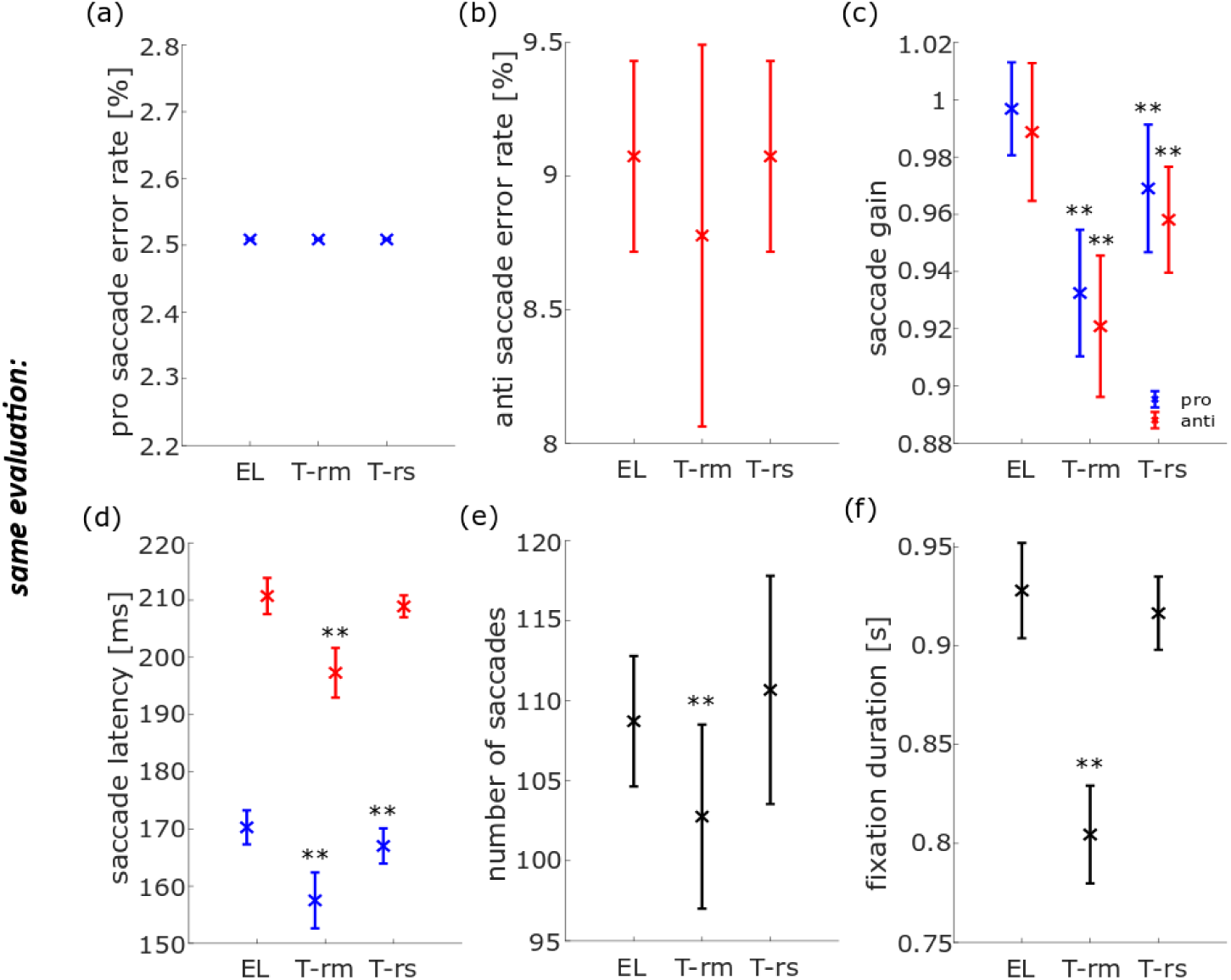
Population data for six eye movement parameters in the pro– and anti-saccade task, measured with the EyeLink 1000 (EL), TOM-research mobile (TOM-rm) and TOM-research stationary (TOM-rs) and analyzed with the same saccade and fixation detectors. The blue data points and their inter-device SD represent pro-saccade data, the red points and their inter-device SD the anti-saccade data and the black points and their inter-device SD the joint pro– and anti-saccade data. (a) Error rate of pro-saccades, (b) error rate of anti-saccades, (c) mean saccade gain, (d) mean saccade latency, (e) mean number of saccades (f) and mean fixation duration. **p < 0.01.

### Free Viewing Task

The spatial accuracy of the TOM-rm was slightly worse than one degree of visual angle, therefore only saccade amplitudes greater than one degree of visual angle were considered in the evaluation of the free-viewing data.

To analyze the eye-movements of the free viewing task we first determined the mean number of fixations per image (stimulus duration: 6s) with the *individual evaluation* approach. We found a significant difference between the EL dataset and the dataset from the TOM-rm concerning the number of fixations (roughly 12 fixations derived from the TOM-rm dataset and 15 fixations derived from the EL-dataset). The values derived from the TOM-rm in general were smaller than those from the EL. To a minor extent, the number of fixations of the TOM-rs was also smaller than that of the EL (t_rm_(20) = 6.63, t_rs_(20) = 2.65, p_rm_ = 5.54 x 10^-6^, p_rs_ = 4.62 x 10^-2^; Figure 6a). Furthermore, we found a significant difference between the EL and the TOM-rm datasets concerning the mean amplitude of saccades, but not between the datasets of the two stationary eye-trackers. Saccade amplitudes as determined from the TOM-rm datasets were on average approx. 15% smaller than those obtained from the EL (t_rm_(20) = 11.06, t_rs_(20) = 1.00, p_rm_ = 1.69 x 10^-9^, p_rs_ = 0.99. Figure 6b). In the mean fixation duration, there was no significant difference between the EL and the TOM-eye-tracker datasets (t_rm_(20) = 0.69, t_rs_(20) = 0.81, p_rm_ = 1.00, p_rs_ = 1.00; Figure 6c).

**Figure 6:**
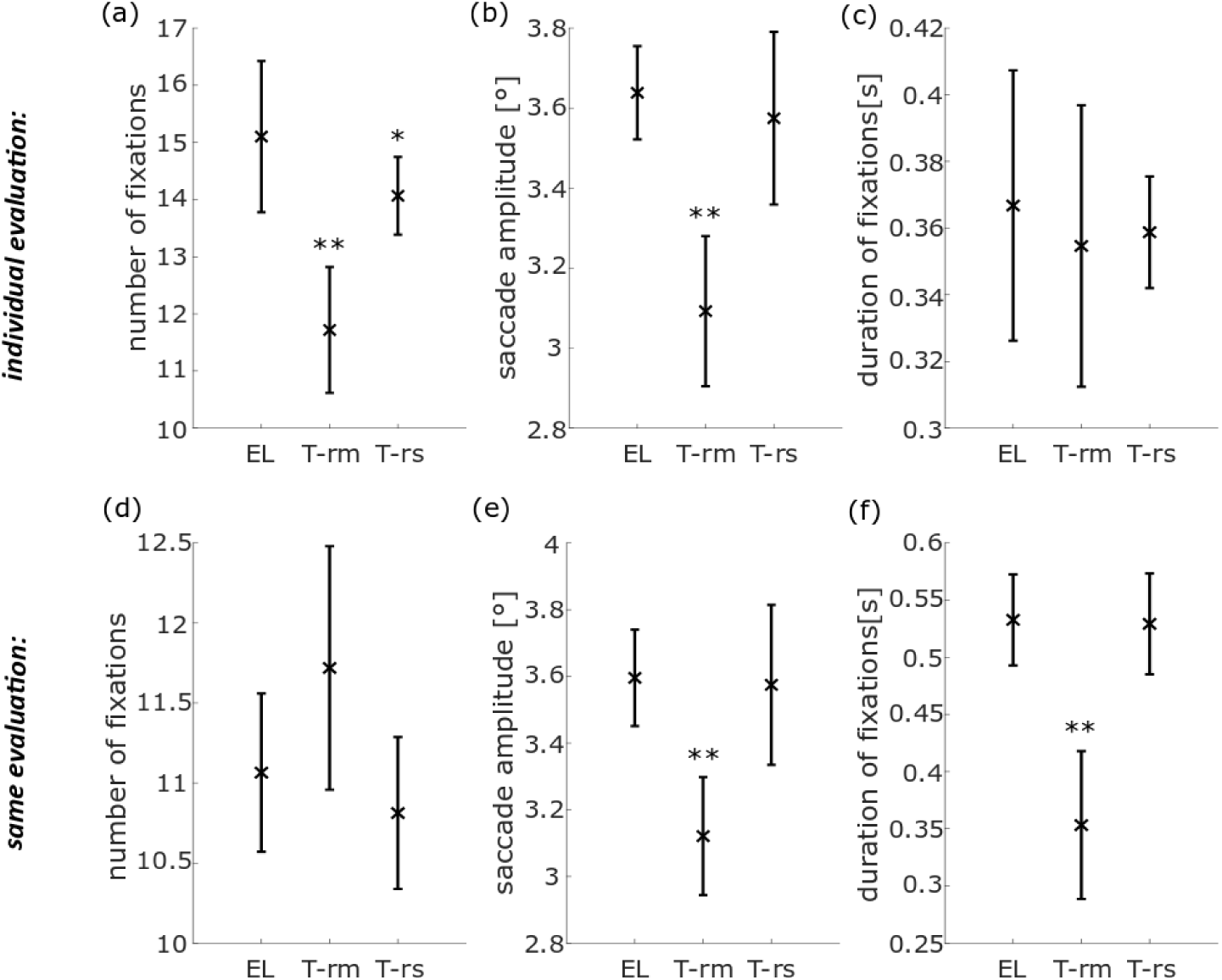
Population data for eye movement parameters in the free viewing task, measured with the EyeLink 1000 (EL), TOM-research mobile (T-rm) and TOM-research stationary (T-rs) and analyzed with the *individual* (a-c) and *same* (e-f) saccade and fixation detector. The cross represents the mean value and the bars their inter-device SD. (a/d) Mean number of fixations per image, (b/e) saccade amplitude and, (c/f) mean fixation duration. *p < 0.05, **p < 0.01.

In a second step, we examined the eye movement parameters in the free viewing task using the *same evaluation* routines to identify potential differences between the three eye-tracking devices. The number of fixations did not differ significantly between the TOM-eye-trackers and the EL (t_rm_(20) = –2.51, t_rs_(20) = 1.93, p_rm_ = 0.06, p_rs_ = 0.20. Figure 6d). The mean saccade amplitude and its inter-device SD are shown in Figure 6e. Just as in the *individual evaluation*, we found a significant difference of saccade amplitudes when applying the *same evaluation* to data from the TOM-rm and the EL (3.6° vs 3.12°), but not to the data form the two stationary eye-trackers (t_rm_(20) = 10.03, t_rs_(20) = 0.27, p_rm_ = 8.99 x 10^-9^, p_rs_ = 1.00). Finally, there was a significant difference between the EL and the TOM-rm dataset concerning the mean fixation duration (0.53s vs 0.35s), but we found no such difference between data from the two stationary eye-trackers (t_rm_(20) = 8.40, t_rs_(20) = 0.30, p_rm_ = 1.63 x 10^-7^, p_rs_ = 1.00. Figure 6f).

The pupillary response, as measured with the EL and the TOM-rs, was evaluated separately for bright (light reflex) and dark images (dark reflex) in the free viewing task. Figure 7 shows the image induced change in normalized pupil area averaged across all subjects over time. Time t = 0s corresponds to the time of displaying the (bright or dark) image. The pupil area was normalized by forming the z-score and this value was set to 0 for t = 0s. The dashed lines indicate the pupil reactions during viewing bright images and the solid lines during viewing dark images. The vertical dashed lines represent the start times of the light reflex (purple: as derived from EL data; cyan: as derived from the TOM-rs data) which were not significantly different (t(20) = –1.02, p = 0.97; Figure 7d & 7e). The times of the dark reflex are represented by the vertical solid purple (EL) and cyan (TOM-rs) lines. No difference could be found here either (t(20) = 0.12, p = 1.00). We also computed the maximum normalized constriction amplitude. Both values did not differ significantly from each other (t(20) = –0.74, p = 1.00. Figure 7b). Finally, the time of maximum constriction as determined from the two datasets was not significantly different (t(20) = –0.45, p = 1.00. Figure 7c).

**Figure 7:**
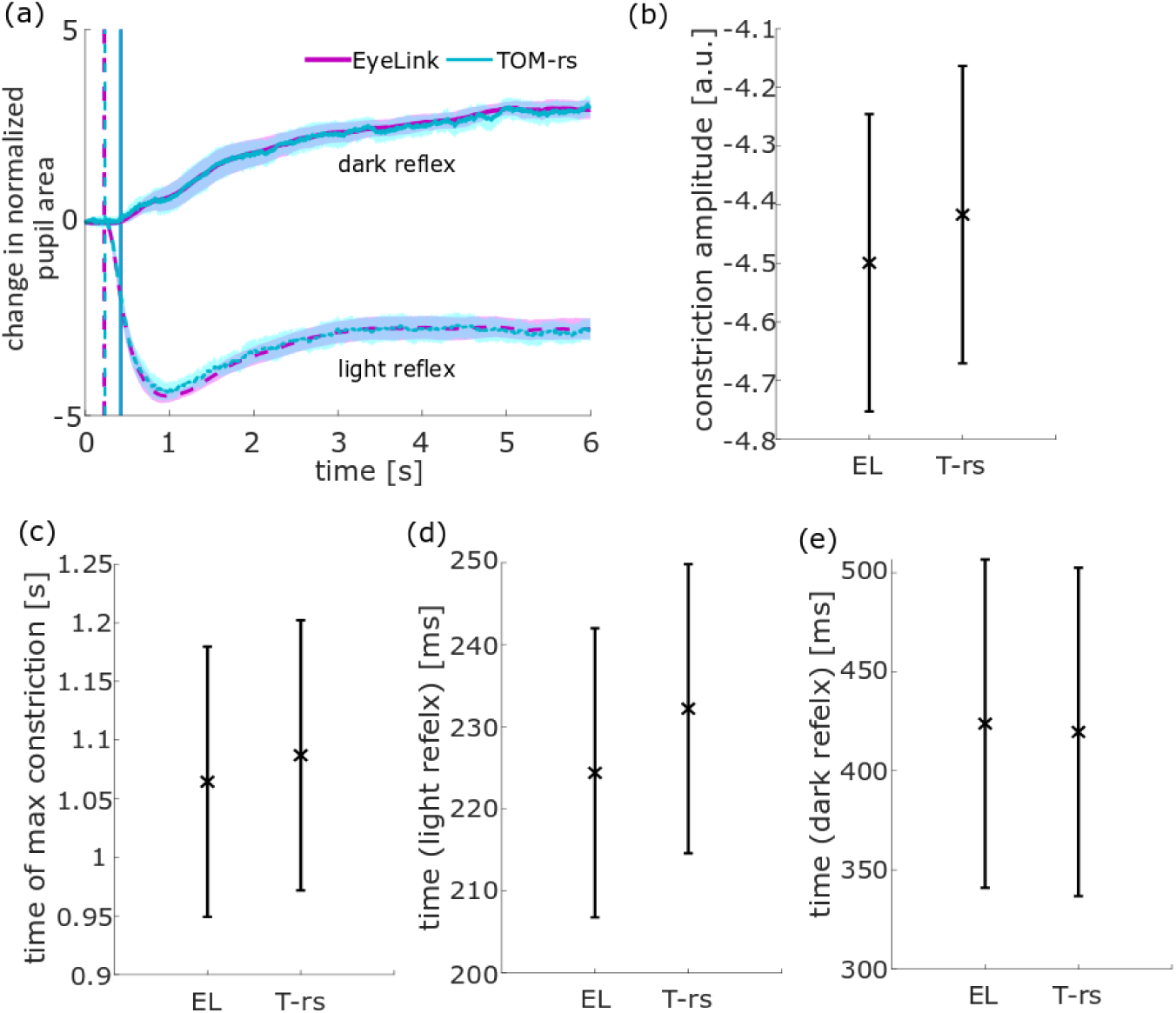
Population data for pupil reaction parameters in the free viewing task, measured with the EyeLink 1000 (EL) and TOM-research stationary (TOM-rs). The error bars represent the inter-device SD. (a) Normalized pupil area averaged over all subjects over time with its 95% confidence interval. The time t = 0s corresponds to the time of displaying the image. The dashed lines show the pupil data from viewing the bright images and the solid lines show the data from viewing the dark images. The vertical dashed purple line represents the start time of the light reflex as determined from the EL-dataset and the vertical dashed cyan line represents the start time as derived from the TOM-rs-dataset. The times of the dark reflex are represented by the vertical purple line (EL) and by the vertical cyan line (TOM-rs). (b) Normalized contraction amplitude of the EL and of the TOM-rs. (c) Time of maximal constriction. (d) Start time of the light reflex and (e) Start time of the dark reflex.

### TOM-rs measurement under ideal IR illumination

The concurrent measurements with all three eye–trackers required a compromise concerning the infrared (IR) illumination (see Methods for details). The EL requires less intense IR illumination. Accordingly, if we had applied the TOM-rs specific illumination, this would have resulted in an overexposure of the EL images which, in turn, would have made concurrent recordings with the EL impossible. Due to the ability of the TOM-rs to flexibly adjust the sampling rate and the shutter speed of the camera, a compromise setting has been employed, in which the IR illumination was close to optimal for the EL, but it was clearly suboptimal for the TOM-rs (the TOM-rm does not require additional IR illumination). Hence, in a final step, we measured a subset of our paradigm (pro– and anti-saccades) with the TOM-rs under ideal IR illumination (TOM-rs ideal). Figure 8 shows the horizontal eye position during a saccade to a peripheral target and back, in the left figure for the compromise setting and in the right figure under the ideal IR-illumination conditions.

**Figure 8:**
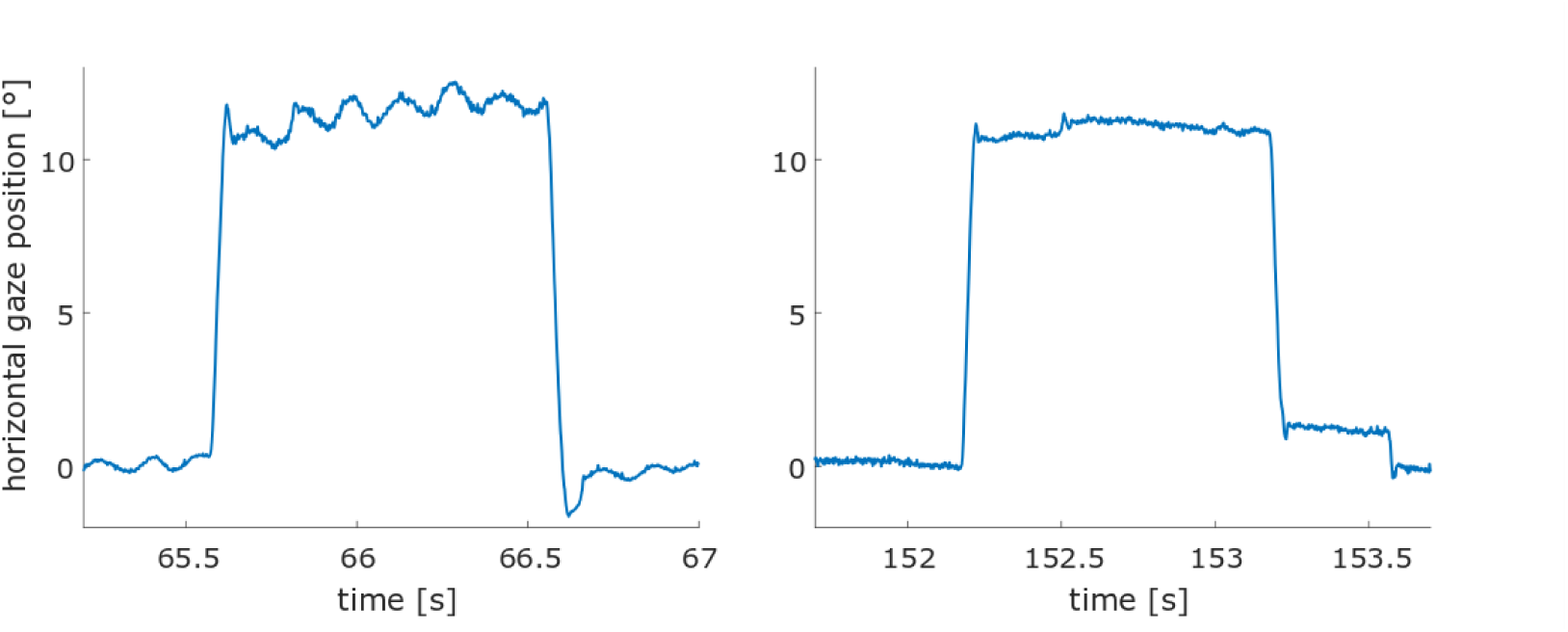
Eye position data over time measured in the pro– and anti-saccade task. Left: Measurement of a saccade to the right under the compromise IR-illumination condition. Right: Measurement of a saccade to the right under the optimized IR-illumination condition.

It becomes obvious that under the compromise IR setting, the noise in the dataset was much higher than under optimized conditions. For a quantitative analysis, we compared the measurements of the TOM-rs under ideal conditions with the previously measured data (parallel measurement with all three eye-trackers) of the EL and the TOM-rs (Figure 9) for the same parameters as before (error rate, gain, latency, number of saccades and fixation duration). No significant difference was found between the measurements under ideal exposure conditions with the TOM-rs and under the compromise condition with the TOM-rs and the EL.

**Figure 9:**
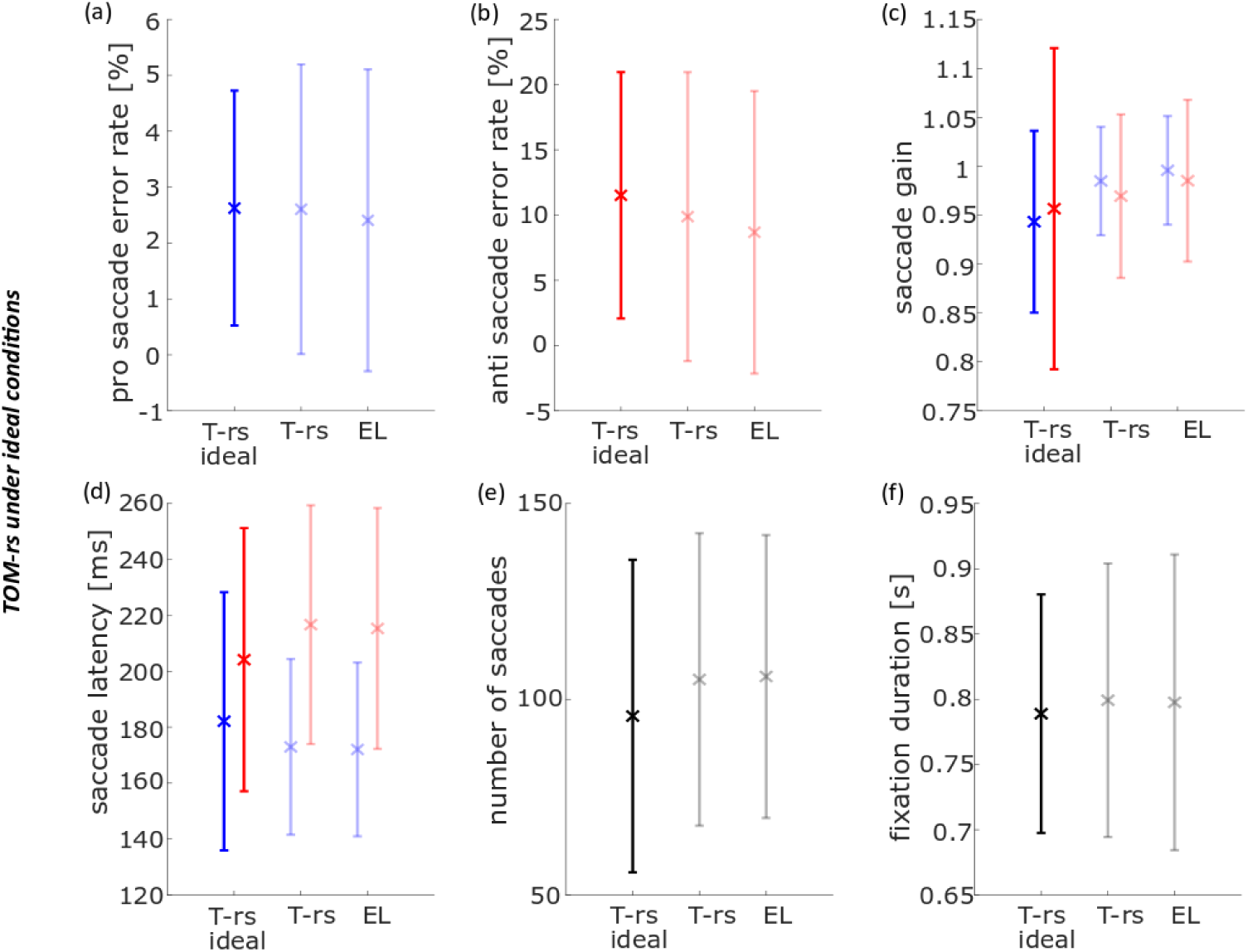
Population data of n = 7 subjects for six eye movement parameters in the pro– and anti-saccade task, measured with the TOM-research stationary under ideal conditions (TOM-rs ideal) and analyzed with individual saccades and fixation detectors, compared with the original EyeLink (EL) and TOM-rs data (transparent). The blue points and their inter-individual SD represent the pro-saccade data, the red points and their inter-individual SD the anti-saccade data and the black points and their inter-individual SD pro– and anti-saccade-data. (a) Error rate of pro-saccades, (b) Error rate of anti-saccades, (c) Mean saccade gain, (d) Mean saccade latency, (e) Mean number of saccades (f) and mean fixation duration.

## Discussion

In this study we quantitatively compared eye-movement parameters as derived from measurements with two novel eye-trackers (TOM-rm and TOM-rs) with those from the EyeLink 1000 (EL). One of these new eye-trackers is a high-resolution stationary eye-tracker (TOM-rs) and the other is a mobile, tablet-based eye-tracker with a frame rate of 30Hz (TOM-rm). The EL is a well-established eye-tracker in oculomotor research and therefore served as a reference system for our study.

### General remarks

In this study, we performed the oculomotor recordings concurrently with all three eye-trackers, which has advantages and disadvantages. One of the advantages was that we were able to compare the identical eye movement parameters quantitatively and thus directly compared the eye-trackers with each other. The disadvantage of this method was that the different eye-trackers have different demands especially concerning the lighting of their environment, so compromises must be made. Recordings with the EL and TOM–rs require IR-light and should take place in a darkened room so that the contrast between pupil and iris increases. Since the TOM-rm requires visible light, our measurement took place in a well-lit room, which can slightly affect the IR spectrum of the EL and TOM-rs Systems (Kunka & Kostek, 2009).

Ideally, the results as obtained from the two stationary eye-trackers (EL and TOM-rs) should not differ significantly neither concerning eye movement data nor results from pupillometry. However, both systems require IR illumination and their IR light sensitivities are different. Our highest priority was to concurrently measure eye-movements with all three systems. Only such an approach allows to compare identical oculomotor data with each other. As a consequence, we had to find a compromise concerning the IR illumination. Due to the fixed focal length lens but high light sensitivity, the required IR illumination for the EL is weaker than for the TOM-rs, which uses a variable focal length zoom lens with slightly lower light sensitivity. Our intention was to keep the measurement conditions between the eye-trackers as similar as possible. Thus, we decided to use an EL-optimal IR illumination, because otherwise the images from the EL would have been overexposed and non-usable. On the other hand, the TOM-rs allows to flexibly adjust the sampling rate and the shutter speed of the camera, which allowed us to find the best setting of this non-optimal illumination. Yet, this compromise still affected the eye-tracking quality of the TOM-rs, since these images tended to be underexposed and the resulting noise could only partially be eliminated by smoothing the data.

For the quantitative comparison, we first evaluated the data of the three eye-trackers with *individually* adapted evaluation software. In this type of evaluation, both hardware-specific and software-specific components were implicitly included in the results. To understand which differences are due to the hardware, we used the *same evaluation* software (that of the TOM-rm) for all three eye-trackers.

### Functional characteristics of the two novel eye-trackers

The TOM-rm is a fully integrated mobile device, which means that a laboratory environment is not necessary for a measurement. Another characteristic of the TOM-rm is that the device is lightweight, easy to use and requires only a short training in handling. Unlike the EL or TOM-rs, the quality of gaze detection depends especially on the lighting condition (in the visible light range). The TOM-rm measures eye-movements at a frame rate of 30Hz, which means that parameters that need a high temporal resolution (e.g., saccade latency) cannot be reliably determined. Figure 10 shows, from a theoretical perspective, the effect of high noise and low sampling frequency in comparison to low noise and high sampling frequency on the fixation and saccade detection. Figure 10 (top) shows that the eye position of the data with the high noise and the low frame rate has the same tendency as the eye position of the data with the low noise and the high frame rate, but under consideration of the single samples, there are huge differences.

**Figure 10:**
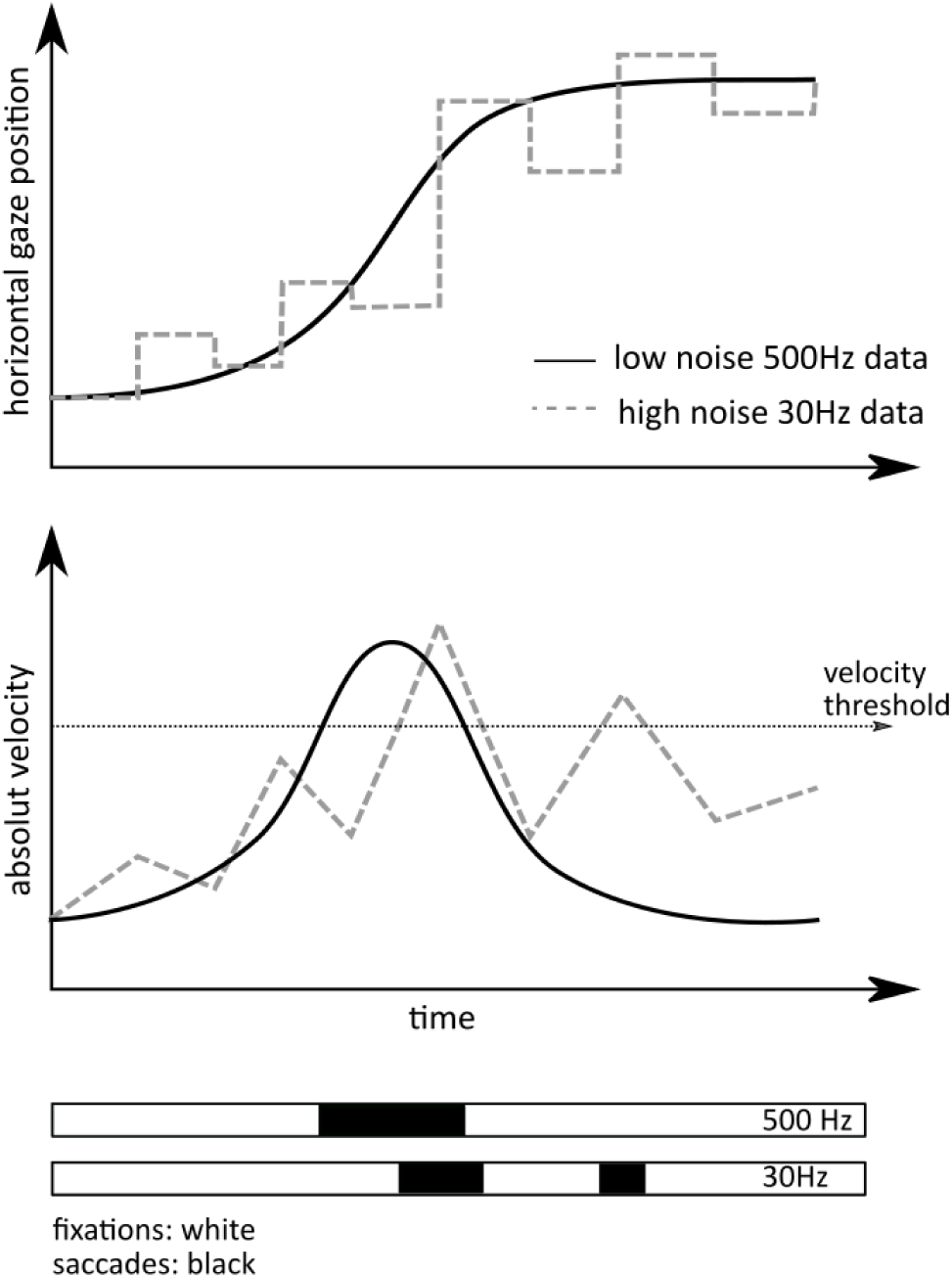
Illustration of the effect of high noise and low frequency on fixations and saccades. According to: Reingold, 2014.

This affects the gaze velocity, resulting in saccades and fixations having a different onset and offset. Most commercially available tablets don’t have a front camera that can detect infrared light, so only visible light can be used for the measurement and the gaze position is detected by capturing the entire iris. The advantage of IR light is that it eliminates unwanted artifacts and leaves unique reflections on the user’s eye (Kunka et al., 2010). In addition, the sclera and iris reflect IR light, while visible light is reflected only by the sclera. Since the sharpest contour is between the iris and pupil and not between the iris and sclera, the pupil can be detected much more precisely with IR light than the iris with visible light (Kunka & Kostek, 2009). In the raw data the noise of the TOM-rm was relatively high as compared to the stationary eye-trackers. Accordingly, it appears challenging to determine oculomotor parameters, which require a high spatial resolution (e.g., saccade gain). Our results confirm this view and suggest that the TOM-rm is suitable for the measurement of oculomotor parameters that don’t require high spatial or temporal resolution, like the pro– and anti-saccade error rate, and measurements that require a flexible setting for the recordings. More accurate measurements, on the other hand, are possible with the TOM-rs. With the high-resolution camera, which can record data with up to 2000Hz, it is possible to perform temporally and spatially accurate and precise measurements. Since the TOM-rs uses a zoom lens with a variable focal length between 16mm-300mm and camera parameters (such as the shutter speed) are freely adjustable, it is rather flexible regarding the measurement environment. Simultaneously, this feature has the disadvantage that the camera settings have to be tuned individually for each environment since errors in the adjustment can significantly degrade data quality.

### Pro– and anti-saccade task

First, we analyzed data from the pro– and anti-saccade task with the parameters being (i) error rate, (ii) saccade gain, (iii) saccade latency (iv), number of saccades and (v) duration of fixation. During an anti-saccade, the subject must suppress a reflexive, visually guided eye movement in the direction of the target and perform a voluntary saccade in the opposite direction. This task is error prone for most subjects, i.e., participants tend to make an erroneous pro-saccade, often followed by a very short latency corrective saccade in the required direction (Coe & Munoz, 2017). This result was confirmed from all three datasets, as derived from the *individual* and the *same evaluations*, respectively (Figures 4a, b and Figures 5a, b). In a comparable anti-saccade gap task, Waldthaler et al. (2021) determined an anti-saccade error rate of almost 20%. In our case, the anti-saccade error rate was lower, i.e., at 9% to 10% on average. Since the anti-saccade error rate increases approximately with 0.5% per year (Mack et al., 2020), this could have been due to the relatively young participants in our study as compared to the study of Waldthaler et al. (2021), in which healthy control subjects were age-matched to patients with Parkinson’s disease, who are usually older. Figure 11(a) shows the horizontal eye position over time is shown for the EL (purple) and the TOM-rs (yellow) during a saccade. The blue symbols represent the start of a saccade and the red the end. The square corresponds to the saccade start/end of the EL and the crosses to the TOM-rs. In the *individual evaluation*, the error rate of the EL was smaller (not significant) than that of the TOM-rs, which was due to the fact that the previously mentioned fast erroneous saccades were not detected by the EL saccade detector in a few cases. Typically, saccades undershoot the target position (Hallett, 1978; Smit, Van Gisbergen, & Cools, 1987; Krappmann, 1998). Our data are in line with these previous results.

**Figure 11:**
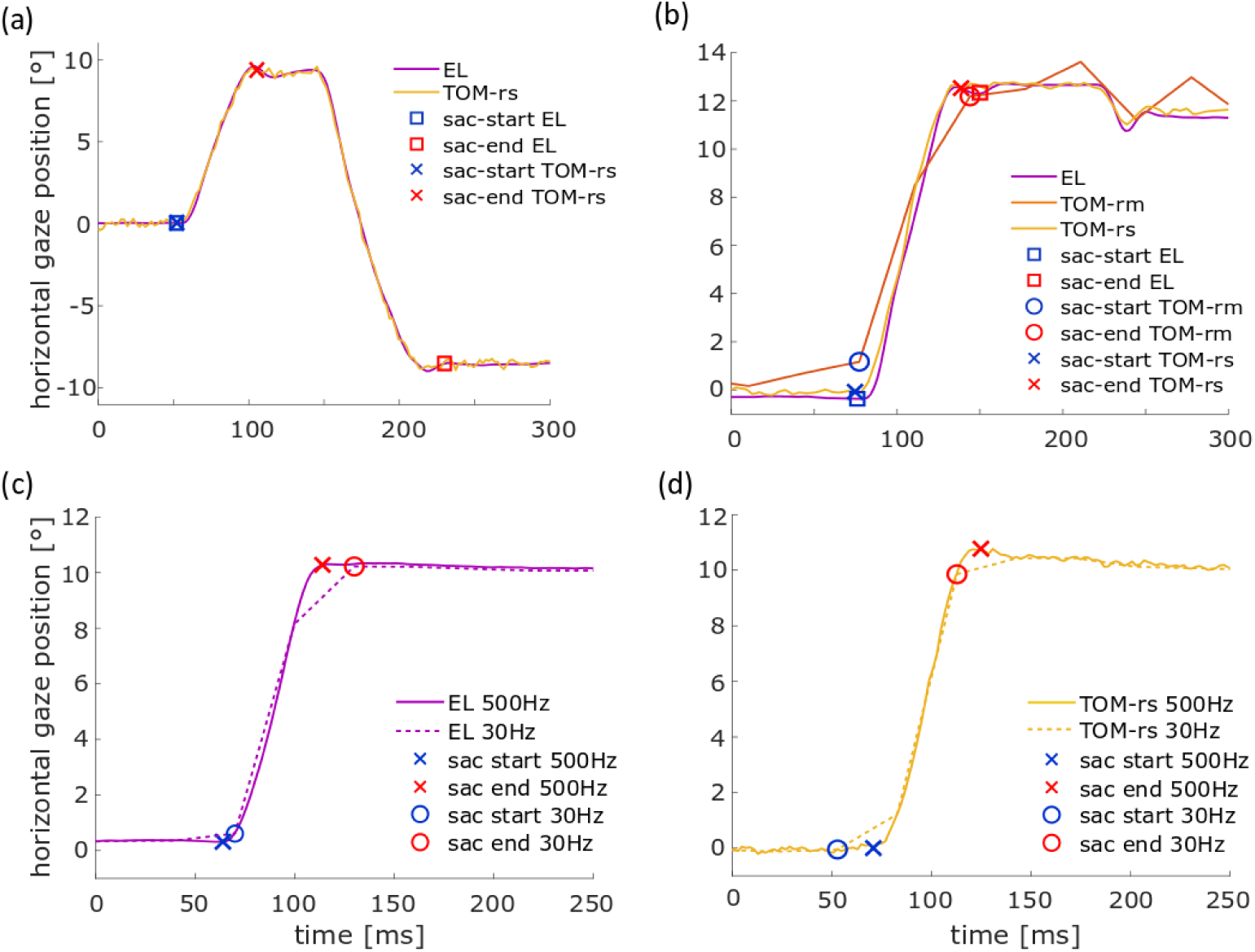
Illustration of the differences in saccade detection in the pro– and anti-saccade task of (a) an erroneous saccade in *individual evaluation* for the eye position data between the standard EyeLink 1000 (EL) and the TOM – research stationary (TOM-rs), (b) the *individual evaluation* of the eye position data between the standard EL, the TOM-rs and the TOM – research mobile (TOM-rm) saccade gain detection, (c)/(d) *individual* (500Hz) *and same* (30Hz) *evaluation of* the eye position data for the EL and the TOM-rs saccade latency. The blue icon represents the saccade start and the red icon the saccade end.

Saccadic gain as derived from the TOM-rm datasets, however, were on average approx. 7% smaller than the respective values derived from the EL datasets. We assume that this difference was related to the lower spatial and temporal resolution, as shown exemplary in Figure 11(b) (TOM Research Mobile). Here the horizontal eye position over time is shown for the EL (purple), the TOM-rm (orange) and the TOM-rs (yellow) during a saccade. The blue symbols represent the start of a saccade and the red the end. The square corresponds to the saccade start/end of the EL, the circles to the TOM-rm and the crosses to the TOM-rs. The saccade of the TOM-rm is smaller than that of the stationary eye-tracker, thus the saccade gain is also smaller.

When evaluating data from all three eye-trackers with the *same evaluation*, saccadic gain as derived from the TOM-rs data was significantly smaller than the value derived from the EL data set. We assume that this was due to the noisier TOM-rs data under non-optimal IR illumination. The gain in the TOM-rs dataset turned out to be reduced only for the *same evaluation*. It appears likely that this seemingly different performance was induced by the down sampling of the dataset. This had a bigger effect on the TOM-rs data due to the compromise in the intensity of the IR illumination than on the EL data. Figure 11(c/d) shows the horizontal eye position over time for the EL (c) and for the TOM-rs (d). The data set with a frame rate of 500Hz is represented by the solid line and the down sampled data (30Hz) by the dashed line. The blue symbols represent the start of a saccade and the red the end. The cross corresponds to the saccade start/end of the EL/TOM-rs at 500Hz and the circles at 30Hz. The compromise reduces the precision, which in turn could reduce the detected saccade amplitude and therefore the saccade gain. The *individual evaluation* shows that the gain as derived from the two stationary eye-trackers was almost the same, but not for the mobile eye-tracker.

For the analysis with the *same evaluation* algorithm, the data first was down-sampled to 30Hz. Anti-saccade latency for all three eye-trackers was higher than for pro-saccades, which corresponds to findings reported in the literature (Everling & Fischer, 1998). The low frame rate and high noise of the TOM-rm results in lower saccade latency than stationary eye-trackers, this effect is shown in Figure 10. Similar to the saccade gain, down sampling to 30Hz affected the latency of the TOM-rs-data more than the EL-data. Figures 11(c)/(d) show that for the down sampled data the saccade start of the TOM-rs was detected earlier than for the non-down sampled data and the EL data.

The number of saccades as derived from the TOM-rm dataset was on average 5% smaller than that from the EL dataset. This difference was mainly a consequence of the saccade duration threshold (saccade duration < 200ms). In Dalmaijer (2014), it was shown that the saccade duration derived from a low-frequency eye-tracker was higher than that from a high-frequency eye-tracker. Nevertheless, we wanted to restrict saccade duration to this upper limit because even larger values appeared non-physiological. On average, (11.00 ± 7.98) saccades per subject were removed for the TOM-rm dataset using this saccade duration criterion. Concerning the *same evaluation* (6.05 ± 5.29) saccades per subject of the EL datasets were removed and (5.24 ± 6.60) saccades per subject of the TOM-rs datasets.

The duration of the fixation in the *same evaluation* was shorter for the TOM-rm than for the two stationary eye-trackers, because small saccades of the stationary eye-trackers in *the individual evaluation* as shown in Figure 12(a) did not interrupt periods of fixations as determined in the *same evaluation* (Figure 12(b)).

**Figure 12:**
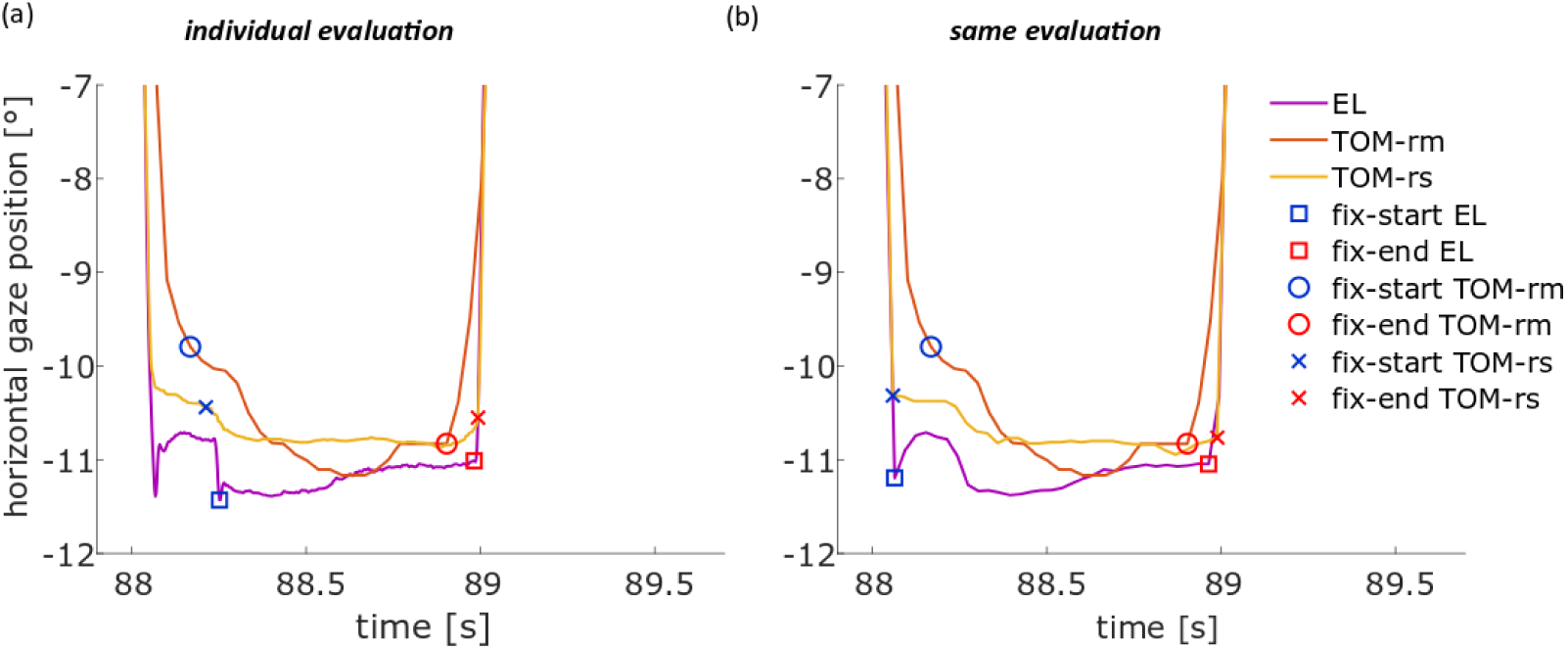
Illustration of the differences in fixation detection for the eye position data between the EyeLink 1000 (EL, blue), TOM – research mobile (TOM-rm, orange) and the TOM – research stationary (TOM-rs, yellow) fixation duration in the pro– and anti-saccade task. The blue square (circle, cross) represents the saccade start and the red square (circle, cross) the saccade end of the EL (TOM-rm, the TOM-rs). (a) one fixation of a representative subject for the *individual evaluation* and (b) one fixation of the same representative subject for the *same evaluation*.

This leads to the fact that the saccades of the stationary eye-trackers in the evaluation with the *same evaluation* were longer than those of the *individual evaluation*. For the *individual evaluation*, no difference could be found between the eye-trackers. If there were no small saccades during fixation, then the saccades of the stationary eye-trackers were longer than those of the TOM-rm, due to the higher frame rate. However, if small saccades interrupt the fixations of the stationary eye-trackers, then these fixations were shorter than those of the TOM-rm. Considering the total number of trials of the subjects, the differences compensate each other and lead to the result that on average there was no difference between the stationary and the mobile eye-trackers.

In a last step we determined all parameters of the pro– and anti-saccade task with the TOM-rs under ideal IR-illumination conditions in seven subjects. As shown in Figure 8, noise was greatly reduced in this optimized condition in contrast to the measurement under compromise conditions. We had assumed that under ideal IR-illumination conditions the saccade gain would be closer to the values previously derived from the EL-dataset. Yet, unexpectedly, this was not the case, as shown in Figure 9(c). Given that these data were recorded from a new cohort of subjects, we speculate that these subjects on average had a lower saccadic accuracy.

### Free viewing task

The investigation of eye-movements during the exploration of images can be a powerful tool for the detection of neurodegenerative diseases. In Parkinson’s disease, for example the saccade amplitudes are on average smaller than in healthy age-matched controls (Matsumoto et al., 2011; Matsumoto et al., 2012). In this study we investigated the performance of the TOM-eye-trackers during a free viewing task with 30 different images (15 bright and 15 dark images).

For both the *individual* and the *same evaluation*, the saccade amplitude as derived from the datasets of the two stationary eye-trackers did not differ from each other. However, the saccade amplitude of the TOM-rm was more than 0.5° (15%) smaller than that of the two stationary eye-tackers (Figure 13(a)). This effect also occurred in the pro– and anti-saccade task.

**Figure 13:**
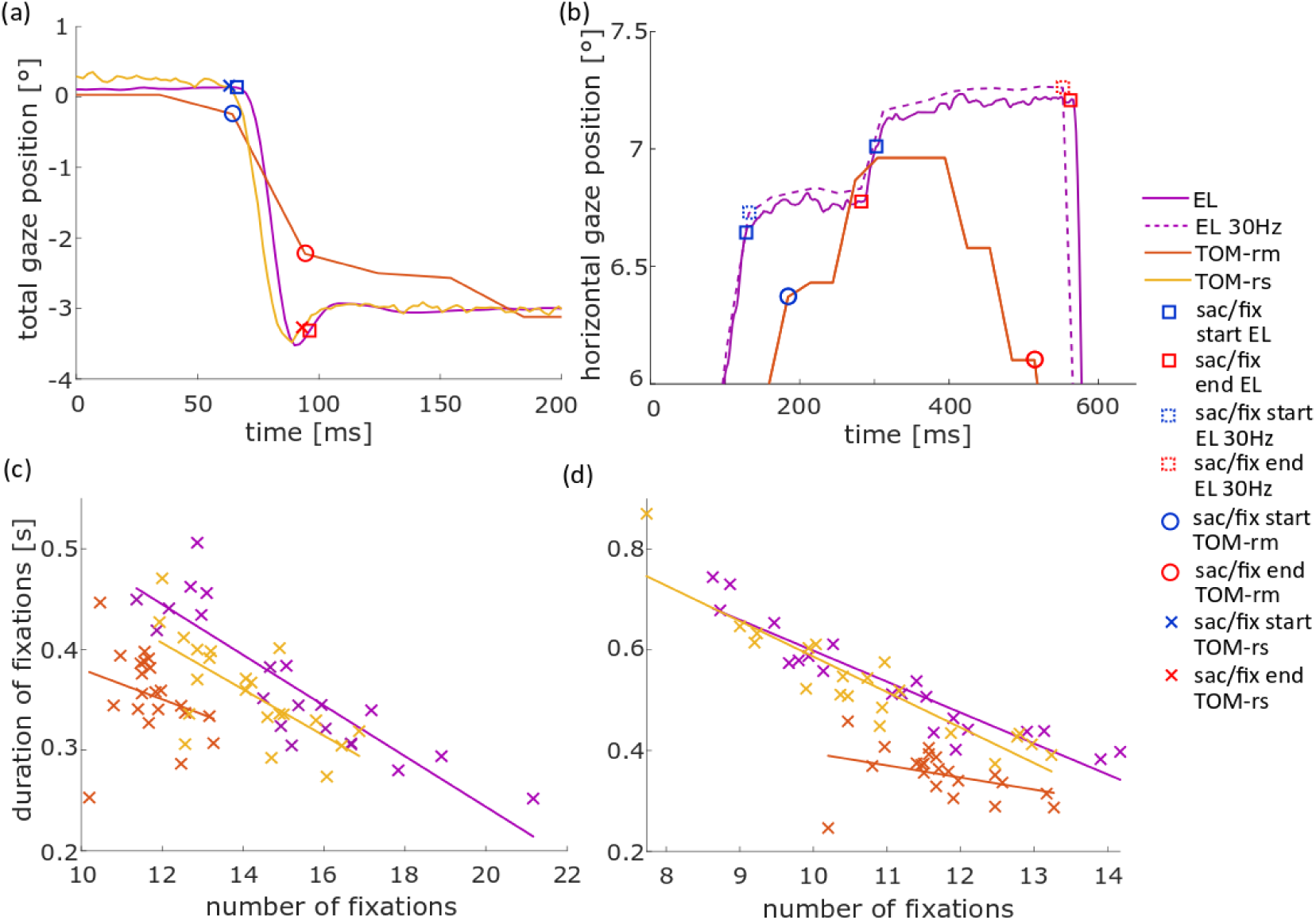
(a) Illustration of the differences in saccade detection for the total gaze position data, representing the combination of vertical and horizontal gaze components, by computing the Euclidian distance of the current gaze position and the center of the screen for each point in time. This results in a comparison between the standard EyeLink 1000 (EL), TOM – research mobile (TOM-rm) and the TOM – research stationary (TOM-rs) saccade amplitude in the free viewing task for the *individual evaluation*. (b) Illustration of the differences in fixation detection for the eye position data between the EL and TOM-rm. (c/d) Illustration of the correlation between the duration of fixations and the number of fixations of the EL, TOM-rm and TOM-rs for the *individual evaluation* (c) and the *same evaluation* (d). The blue symbols (square, circle, cross) represent the saccade/fixation start and the red symbols the end of a saccade.

When we determined the fixation duration with the *same evaluation*, the duration was lower for the EL and TOM-rs than for the *individual evaluation*. This was mainly since the spatial range (t1 and t2) for the fixation detector of the TOM-rm was chosen relatively high (see Methods for details). This range was obviously too high for the TOM-rs and EL datasets and resulted in fixations that were interrupted by small saccades, which however were not detected separately. Accordingly, the *individual evaluation* was better suited for the two stationary eye-trackers. If we compare the TOM-rs with the EL for the evaluation with the *individual* and *same* fixation detector, we found no difference between the eye-trackers. Values derived from the TOM-rm did not differ from those from the two stationary eye-trackers concerning fixation duration in the *individual evaluation*, but for the *same evaluation*. This also suggests that the fixation detector in the *same evaluation* was not suitable for the stationary devices.

For a given temporal interval, the number of fixations is inversely proportional to the fixation duration. If the number of detected fixations increases, the duration of the detected fixations must decrease. This is implicitly reflected in the data shown in Figure 13(c/d).

In the *same evaluation* we found a significant difference between the TOM-rm and the stationary eye-tracker data. Like in the pro– and anti-saccade task, the main reason was that interruptions due to small saccades could not be detected because of the higher noise in the TOM-rm data.

In addition, we also found a small difference between the TOM-rm and the EL in the *individual evaluation*. Due to the compromise conditions, shorter fixations cannot always be reliably detected.

Not only eye-movements but also the pupillary response can be an important biomarker for neurological diseases, as shown in Wang et al. (2016). In this study we investigated the difference in pupillary response between the two stationary eye-trackers while viewing bright and dark images. According to the literature, the latency of the constriction onset of the light reflex is in the range of 230 to 357ms and about 445ms for the dark reflex (Bergamin & Kardon, 2003; Wang et al., 2018). The data of our current study for the light and dark reflex showed similar values. In the light and dark reflex, the EL and the TOM-rs datasets did not differ significantly. The time-course of the pupil areas while viewing the bright images of the two eye-trackers did not show any difference, as was the case for the contraction amplitude for the TOM-rs. However, the constriction amplitude of the TOM-rs was lower, however not significant, than of the EL. Like for the eye movement tasks, the differences probably were caused by the compromise in IR illumination. Without optimal illumination the noise increases and pupil detection becomes less reliable.

## Conclusion

We conclude that simultaneous measurement with three eye-trackers with different demands on the environment, requires compromises that affect the data quality of the eye-trackers. Nevertheless, this compromise is worth it, because it allows a direct and quantitative comparison between the eye-trackers. Our results show that the mobile tablet-based eye-tracker (TOM-rm) is suitable for certain basic oculomotor experiments like the pro– and anti-saccade task. Experiments in which parameters are collected that require a high frame rate or a high spatial accuracy or precision, such as saccade-latency or gain require eye tracking systems with a high spatial and temporal resolution. Our results show that – given ideal IR illumination – the quality of oculomotor data obtained from the TOM-rs are *on a par* with those obtained from a reference in the field, i.e., the EyeLink 1000.

## Abbreviations

TOM-rm: Thomas-Oculus-Motus – research mobile
TOM-rs: Thomas-Oculus-Motus – research stationary
EL: EyeLink 1000 Plus

## Acknowledgments

This work was supported by the German Federal Ministry of Education and Research (BMBF), Project DIADEM and by Hessisches Ministerium für Wissenschaft und Kunst, HMWK (Clusterproject The adaptive Mind – TAM).

The study was planned by A.K., S.D. and F.B., A.K. and research assistants conducted the experiment after instruction and training by A.K.. A.K. performed the analyses of the data. The manuscript was written by A.K., S.D. and F.B.. The two TOM eye-trackers are developed and distributed by Thomas RECORDING GmbH (Giessen, Germany). S.D. was involved in the development of the two TOM eye-trackers as the chief scientific officer of Thomas RECORDING and has been paid by Thomas RECORDING during that time. As part of the collaborative research project DIADEM the eye-trackers were freely provided to the Dept. Neurophysics (Philipps-Universität Marburg, Germany).

## Data Availability Statement

The raw data supporting the conclusions of this article will be made available by the authors, without undue reservation.

## Supplementary Material

**Table S1:** Inter-individual mean values. Inter-individual SD (interi. SD) and inter-device SD (interd. SD) for eye movement parameters in the free viewing task. measured with the EyeLink 1000 (EL). TOM-research mobile (TOM-rm) and TOM-research stationary (TOM-rs) and analyzed with individual saccades and fixation detector (top) and with the same saccades and fixation detector (bottom). The boldly highlighted values differ significantly from the EL values.

| *Pro- & Anti-Saccades task: individual evaluation* |  |  |  |  |  |  |  |  |  |
| --- | --- | --- | --- | --- | --- | --- | --- | --- | --- |
|  | EL |  |  | TOM-rm |  |  | TOM-rs |  |  |
|  | mean value | interi. SD | interd. SD | mean value | interi. SD | interd. SD | mean value | interi. SD | interd. SD |
| *ER pro [%]* | 2.41 | 2.70 | 0.43 | 2.60 | 2.59 | 0.22 | 2.60 | 2.59 | 0.22 |
| *ER anti [%]* | 8.68 | 10.83 | 1.53 | 9.32 | 10.93 | 1.04 | 9.88 | 11.07 | 1.52 |
| *Num. sac.* | 105.81 | 36.13 | 3.64 | 100.14 | 36.21 | 5.75 | 105.05 | 37.35 | 3.63 |
| *Dur. of fix. [s]* | 0.80 | 0.11 | 0.03 | 0.80 | 0.08 | 0.05 | 0.80 | 0.10 | 0.04 |
| *Gain pro* | 1.00 | 0.06 | 0.02 | **0.93** | **0.06** | **0.03** | 0.98 | 0.06 | 0.03 |
| *Gain anti* | 0.99 | 0.08 | 0.03 | **0.92** | **0.08** | **0.02** | 0.97 | 0.08 | 0.03 |
| *Lat. pro [ms]* | 172.04 | 31.08 | 2.39 | **155.76** | **28.51** | **3.56** | 172.94 | 31.42 | 2.19 |
| *Lat. anti [ms]* | 215.32 | 43.07 | 2.97 | **197.09** | **40.23** | **4.95** | 216.67 | 42.69 | 2.52 |
| *Pro- & Anti-Saccades task: same evaluation* |  |  |  |  |  |  |  |  |  |
|  | EL |  |  | TOM-rm |  |  | TOM-rs |  |  |
|  | mean value | interi. SD | interd. SD | mean value | interi. SD | interd. SD | mean value | interi. SD | interd. SD |
| *ER pro [%]* | 2.51 | 2.66 | 0.00 | 2.51 | 2.66 | 0.00 | 2.51 | 2.66 | 0.00 |
| *ER anti [%]* | 9.05 | 8.90 | 0.36 | 8.75 | 8.85 | 0.71 | 9.05 | 8.90 | 0.36 |
| *Num. sac.* | 108.71 | 38.19 | 4.07 | **102.76** | **38.30** | **5.75** | 110.67 | 39.37 | 7.12 |
| *Dur. of fix. [s]* | 0.93 | 0.14 | 0.03 | **0.80** | **0.08** | **0.05** | 0.92 | 0.13 | 0.03 |
| *Gain pro* | 1.00 | 0.05 | 0.02 | **0.93** | **0.06** | **0.02** | **0.97** | **0.06** | **0.02** |
| *Gain anti* | 0.99 | 0.08 | 0.02 | **0.92** | **0.07** | **0.02** | **0.96** | **0.08** | **0.02** |
| *Lat. pro [ms]* | 170.34 | 32.80 | 2.96 | **157.38** | **29.09** | **4.83** | **167.16** | **32.30** | **2.99** |
| *Lat. anti [ms]* | 210.74 | 42.36 | 3.19 | **197.29** | **39.64** | **4.44** | 209.04 | 41.27 | 1.97 |

**Table S2:**
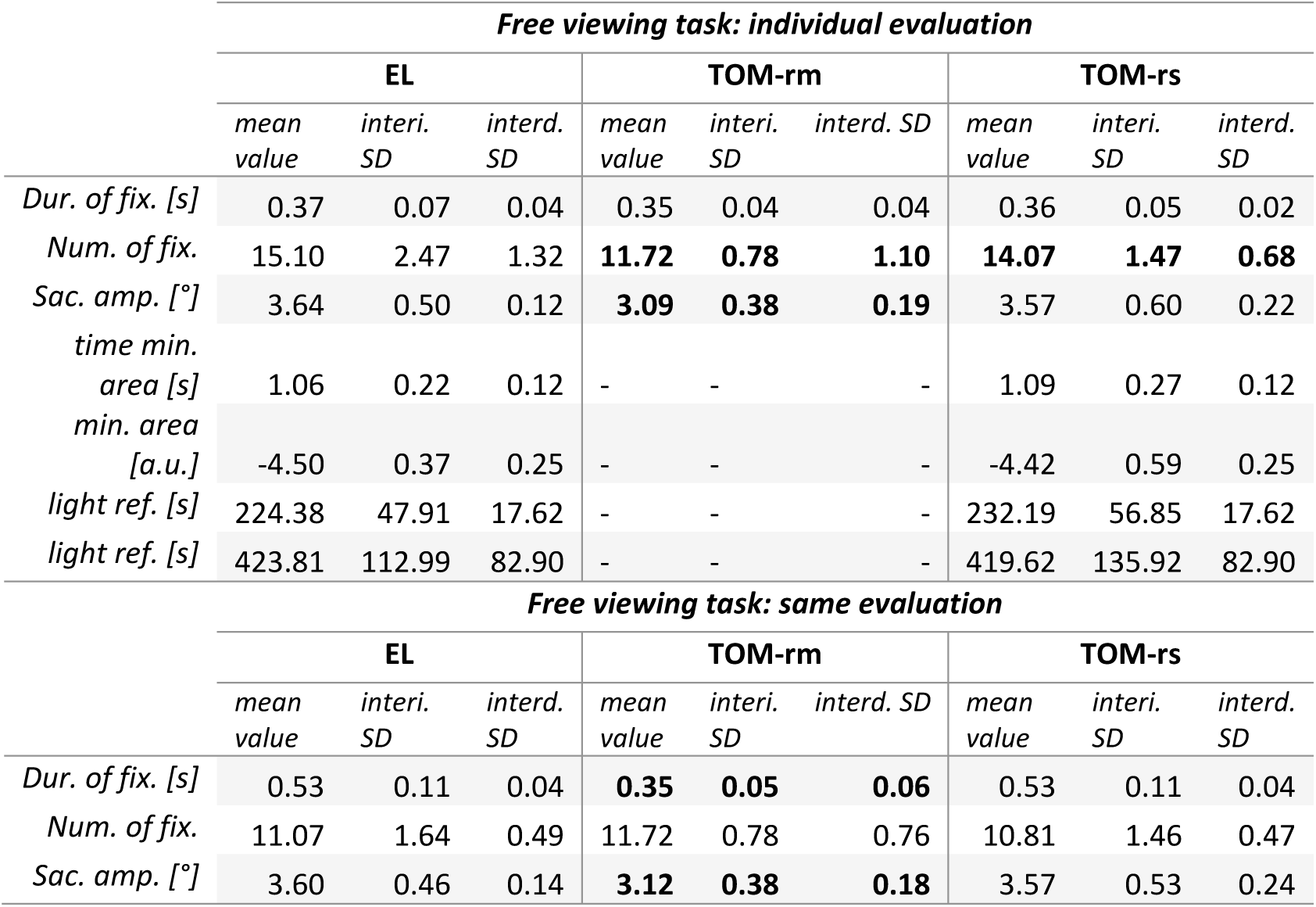
Inter-individual mean values, inter-individual SD (interi. SD) and inter-device SD (interd. SD) for eye movement parameters in the free viewing task. measured with the EyeLink 1000 (EL). TOM-research mobile (TOM-rm) and TOM-research stationary (TOM-rs) and analyzed with individual saccades and fixation detector (top) and with the same saccades and fixation detector (bottom). The boldly highlighted values differ significantly from the EL values.

